# Antibody repertoire induced by SARS-CoV-2 spike protein immunogens

**DOI:** 10.1101/2020.05.12.091918

**Authors:** Supriya Ravichandran, Elizabeth M. Coyle, Laura Klenow, Juanjie Tang, Gabrielle Grubbs, Shufeng Liu, Tony Wang, Hana Golding, Surender Khurana

**Affiliations:** Division of Viral Products, Center for Biologics Evaluation and Research (CBER), FDA, Silver Spring, Maryland, USA, 20871

**Keywords:** SARS-CoV-2, Vaccine, Spike, Neutralization, Epitope, Virus, Antibody, Antigen

## Abstract

Multiple vaccine candidates against SARS-CoV-2 based on viral spike protein are under development. However, there is limited information on the quality of antibody response generated following vaccination by these vaccine modalities. To better understand antibody response induced by spike protein-based vaccines, we immunized rabbits with various SARS-CoV-2 spike protein antigens: S-ectodomain (S1+S2) (aa 16-1213), which lacks the cytoplasmic and transmembrane domains (CT-TM), the S1 domain (aa 16-685), the receptor-binding domain (RBD) (aa 319-541), and the S2 domain (aa 686-1213 as control). Antibody response was analyzed by ELISA, Surface Plasmon Resonance (SPR) against different Spike proteins in native conformation, and a pseudovirion neutralization assay to measure the quality and function of the antibodies elicited by the different Spike antigens. All three antigens (S1+S2 ectodomain, S1 domain, and RBD) generated strong neutralizing antibodies against SARS-CoV-2. Vaccination induced antibody repertoire was analyzed by SARS-CoV-2 spike Genome Fragment Phage Display Libraries (SARS-CoV-2 GFPDL), which identified immunodominant epitopes in the S1, S1-RBD and S2 domains. Furthermore, these analyses demonstrated that surprisingly the RBD immunogen elicited a higher antibody titer with 5-fold higher affinity antibodies to native spike antigens compared with other spike antigens. These findings may help guide rational vaccine design and facilitate development and evaluation of effective therapeutics and vaccines against COVID-19 disease.

**One Sentence Summary:** SARS-CoV-2 Spike induced immune response

## INTRODUCTION

The ongoing pandemic of SARS-CoV-2 has resulted in more than 2 million human cases and 125,000 deaths as of 15^th^ April 2020. Therefore, development of effective vaccines for prevention and medical countermeasures for treatment of SARS-CoV-2 infection is a high global priority. The spike glycoprotein has been identified as the key target for protective antibodies against both SARS-CoV-1 and SARS-CoV-2(*1-4*). Consequently, multiple versions of the SARS-CoV-2 spike proteins are currently under evaluation as vaccine candidates utilizing different modalities and delivery systems(*5*). However, only limited knowledge exists on antibody repertoire or quality of the immune response generated following vaccination by different spike vaccine antigens. Therefore, it is important to perform comprehensive evaluation of post-vaccination antibody response to elucidate the quality of the immune responses elicited by spike-based vaccine candidates to determine immune markers that may predict clinical benefit which can facilitate evaluation of vaccine candidates.

To better understand vaccination-induced antibody response, we immunized rabbits with several SARS-CoV-2 spike proteins: the S-ectodomain (S1+S2) (aa 16-1213) lacking the cytoplasmic and transmembrane domains (delta CT-TM), the S1 domain (aa 16-685), the receptor-binding domain (RBD) (aa 319-541), and the S2 domain (aa 686-1213), as a control. Post-vaccination sera were analyzed by Genome Fragment Phage Display Libraries covering the entire spike gene (SARS-CoV-2 GFPDL) to determine the polyclonal antibody epitope repertoire generated following vaccination as previously applied for other diseases(*6-10*). In addition, we employed several antibody binding assays (ELISA, Surface Plasmon Resonance (SPR) based real-time kinetics assay) (*10-12*) and an *in vitro* SARS-CoV-2 pseudovirion neutralization assay to measure the quality and function of the antibodies elicited by the different SARS-CoV-2 spike antigens. This study could inform development and evaluation of SARS-CoV-2 vaccines and therapeutics based on the spike glycoprotein.

## RESULTS

### Rabbit immunization with SARS-CoV-2 Spike antigens

Most spike-based vaccines currently under development are designed to contain the receptor-binding domain (RBD; aa 319-541) in some form. Therefore, we evaluated four different commercially available SARS-CoV-2 spike protein and subdomains: the Spike S1+S2 ectodomain (aa 16-1213), the S1 domain (aa 16-685), RBD domain (aa 319-541), and the S2 domain (aa 686-1213) as a control, which is devoid of RBD (Fig. 1A, Suppl. Fig. 1). Theese spike proteins were either produced in HEK 293 mammalian cells (S1 and RBD) or insect cells (S1+S2 ectodomain and S2 domain). The purified S1+S2 ectodomain, the S1 domain, and the RBD proteins retained the functional activity as demonstrated in SPR assay using human ACE2 protein, the SARS-CoV-2 receptor (Fig. 1B). The S1+S2 ectodomain, S1 domain and RBD (black, blue and red binding curves, respectively) demonstrated high-affinity interaction with human ACE2. The control S2 domain protein (green curve), lacking the RBD, did not bind to human ACE2, demonstrating specificity of this receptor-binding assay (Fig.1B).

**Figure 1:**
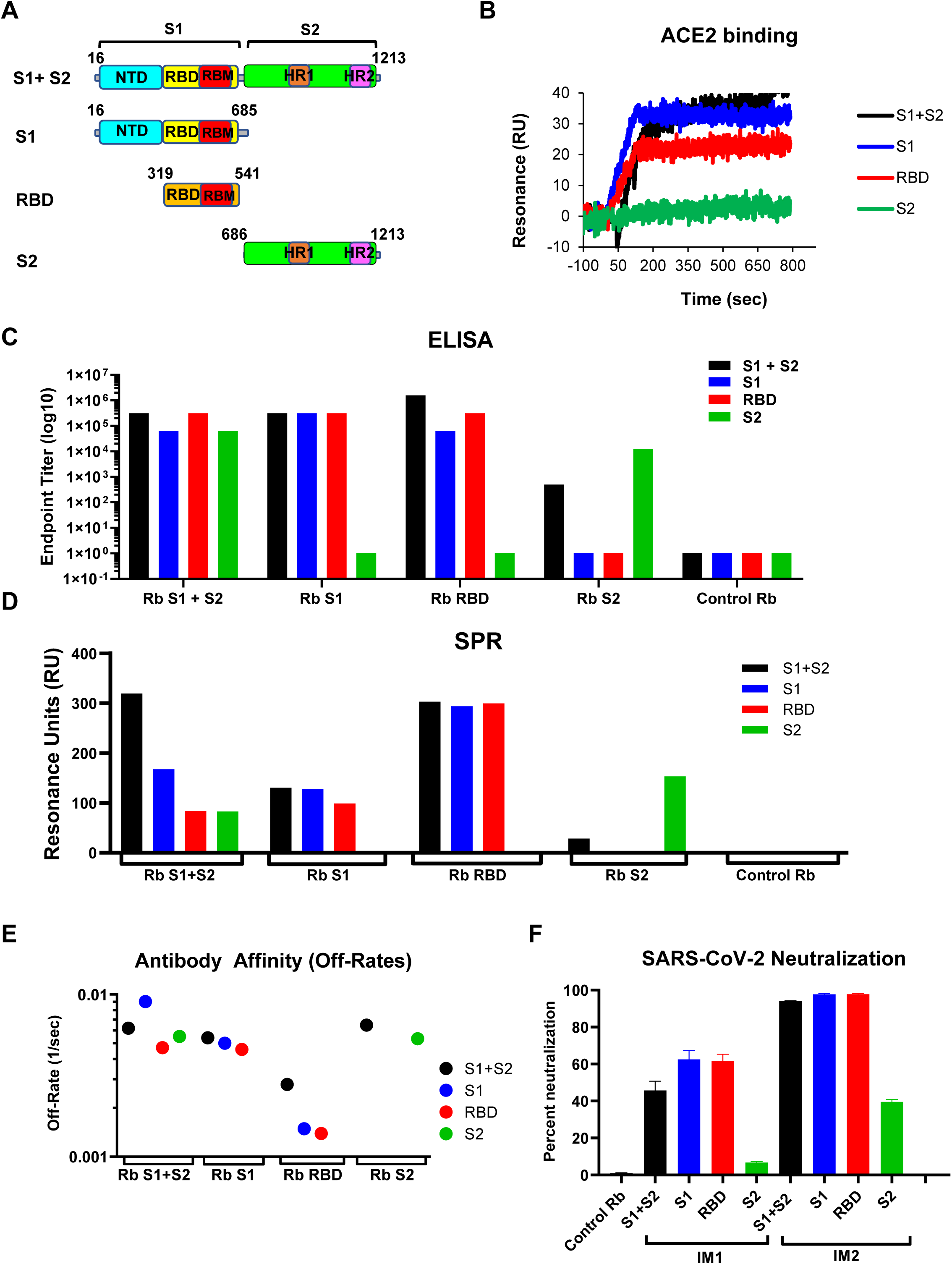
SARS-CoV-2 spike binding and SARS-CoV-2 neutralization by serum antibodies generated following rabbit immunization with spike antigens. A) Schematic representation of the SARS-CoV-2 spike protein and subdomains. Spike S1+S2 ectodomain (aa 16-1213) lacks the cytoplasmic and transmembrane domains (CT-TM), S1 domain (aa 16-685), RBD domain (aa 319-541), and S2 domain (aa 686-1213), all containing 6x His tag at C-terminus, were produced in either HEK 293 mammalian cells (S1 and RBD) or insect cells (S1+S2 ectodomain and S2 domain). (B) Binding of purified proteins to human ACE2 proteins in SPR. Sensorgrams represent binding of purified spike proteins on His-captured chips to 5 µg/mL human ACE2 protein. (C) Anti-spike reactivity of post-immunization rabbit sera. Serial dilutions of post-second vaccination rabbit sera were evaluated for binding to various spike proteins and domains (S1+S2; black, S1; blue, RBD; red, and S2; green) in ELISA. Representative titration curves are shown in Fig. S2. End-point titers of the serum samples were determined as the reciprocal of the highest dilution providing an optical density (OD) twice that of the negative control (no serum was used as negative control). (D) SPR binding of antibodies from rabbits immunized twice with SARS-CoV-2 antigens to spike protein and domains from SARS-CoV-2 (S1+S2; black, S1; blue, RBD; red, and S2; green). Total antibody binding is represented in resonance units in this figure for 10-fold serum dilution. All ELISA and SPR experiments were performed twice and the researchers performing the assay were blinded to sample identity. The variations for duplicate runs of ELISA and SPR were <8% and <5%, respectively. The data shown are average values of two experimental runs. (E) Antibody off-rate constants, which describe the fraction of antigen–antibody complexes that decay per second, were determined directly from the serum/ sample interaction with SARS-CoV-2 spike ectodomain (S1+S2), S1, S2, and RBD using SPR in the dissociation phase only for the sensorgrams with Max RU in the range of 20–100 RU. (F) Virus neutralization titers were measured against SARS-CoV-2-FBLuc in a single-round pseudovirus neutralization assay in triplicates (see Methods). The average percentage inhibition after the first and second vaccination (1:40 serum dilution) for each group are shown. Pre-vaccination rabbit sera also did not neutralize in PsVN assay (Control Rb).

Female New Zealand white rabbits were immunized twice intra-muscularly at a 14-day interval with 50 μg of the purified proteins mixed with Emulsigen Adjuvant. Sera were collected before (pre-vaccination) and after the first and second vaccination and analyzed for binding antibodies in ELISA and SPR, in a pseudovirion neutralization assay, and by GFPDL analysis.

### Antibody Response following immunization with different Spike antigens

Serial dilutions of post-second vaccination rabbit sera were evaluated for binding of serum IgG to various spike proteins and domains in ELISA (S1+S2; black, S1; blue, RBD; red, and S2; green) (Fig. 1C). Representative titration curves to spike ectodomain (S1+S2) and to the RBD in IgG-ELISA are shown in Suppl. Fig. 2. End-point titers of the serum IgG were determined as the reciprocal of the highest dilution providing an optical density (OD) twice that of the negative control (Fig. 1C). All four immunogens elicited strong IgG binding to the spike ectodomain (S1+S2). Binding to the individual domains (S1, S2, and RBD) was specific, in that sera generated by S2 vaccination bound to S2, but not to S1 or RBD, and vice-versa (Fig. 1C).

SPR allows following antibody binding to captured antigens in real-time kinetics, including total antibody binding in resonance units (Max RU) and affinity kinetics (Suppl. Fig. 3). In ELISA, the antigens directly coated in the wells can be partially denatured increasing the likelihood of presenting epitopes that are not seen on the native form of the protein by the polyclonal serum IgG. On the other hand, in our SPR, the purified recombinant spike proteins were captured to a Ni-NTA sensor chip to maintain the native conformation (as determined by ACE2 binding) to allow comparisons of binding to and dissociation from the four proteins. Importantly, the protein density captured on the chip surface is low (200 RU) and was optimized to measure primarily monovalent interactions, so as to measure the average affinity of antibody binding in the polyclonal serum (*8, 13*). Additionally, while ELISA measured only IgG binding, in SPR, all antibody isotypes contributed to antibody binding to the captured spike antigen. In the current study, all rabbit sera contained anti spike antibodies that were at least 86% IgG (data not shown). Serial dilutions of post-vaccination serum were analyzed for binding kinetics with different spike proteins (Suppl. Fig. 3). The spike ectodomain (S1+S2) generated antibodies that predominantly bound to S1+S2 (black bar), followed by the S1 protein (blue bar), and 3-fold lower antibody binding to the RBD and the S2 domain (red and green bars, respectively) (Fig. 1D). The S1 domain antigen induced antibodies that bound with similar titers (Max RU values) to the S1+S2, S1 and RBD proteins (black, blue and red bars, respectively), and did not show reactivity to the S2 domain (green bar). However, the antibody reactivity of rabbit anti-S1 serum to S1+S2 domain was 3-fold lower than the antibodies in the rabbit anti-S1+S2 serum. RBD immunization generated similar high-titer antibody binding to S1+S2, S1 and RBD (black, blue and red bars, respectively), (Fig. 1D). In contrast, the S2 domain induced antibodies that primarily bound to homologous S2 antigen (green bars) and only weakly binding to the S1+S2 ectodomain (black bars), and no binding to either S1 or RBD (Fig. 1D).

Antibody off-rate constants, which describe the fraction of antigen–antibody complexes that decay per second, were determined directly from the serum sample interaction with SARS-CoV-2 spike ectodomain (S1+S2), S1, S2, and RBD using SPR in the dissociation phase only for sensorgrams with Max RU in the range of 20–100 RU (Suppl. Fig. 3) and calculated using the BioRad ProteOn manager software for the heterogeneous sample model as described before(*11*). These off rates provide additional important information on the affinity of the antibodies following vaccination with the different spike proteins that are likely to have an impact on the antibody function *in vivo*, as was observed previously in studies with influenza virus, RSV and Ebola virus (*13-15*). Surprisingly, we observed significant differences in the affinities of antibodies elicited by the four spike antigens (Fig. 1E). Specifically, the RBD induced 5-fold higher affinity antibodies (slower dissociation rates) against S1+S2 (black), S1 (blue) and RBD (red) proteins, compared with the post-vaccination antibodies generated by other three immunogens (Fig. 1E).

SARS-CoV-2 neutralization was measured using SARS-CoV-2-FBLuc in a single-cycle PsVN assay in Vero E6 cells. The average percent inhibition by post-first and post-second rabbit vaccination are shown in Fig. 1F. Pre-vaccination rabbit sera (Control Rb) did not neutralize SARS-CoV-2 in PsVN assay. Sera generated by S1+S2-ectodomain, S1 and RBD (1:40 dilution) (but not anti-S2) showed 50-60% virus neutralization after a single vaccination, and 93-98% virus inhibition by the post-second vaccination sera (Fig. 1F).

### Epitope repertoires recognized by antibodies generated against SARS-CoV-2 spike antigens

The constructed SARS-CoV-2 GFPDL contains sequences ranging from 50-1500 bp long from the spike gene (GenBank #MN908947) with >10^7.2^ unique phage clones. The SARS-CoV-2-GFPDL displayed linear and conformational epitopes with random distribution of size and sequence of inserts that spanned the entire spike gene. SARS-CoV-2 GFPDL panning with individual post-second vaccination rabbit sera were conducted as described in Methods. The numbers of IgG-bound SARS-CoV-2 GFPDL phage clones with different serum sample ranged between 2.6 × 10^4^ to 9.8 × 10^5^/mL (Fig. 2A). Graphical distribution of representative clones with a frequency of ≥2, obtained after affinity selection, and their alignment to the spike protein of SARS-CoV-2 are shown for the four vaccine groups (Fig. 2 B-E). The spike (S1+S2) ectodomain induced diverse antibody response that included strong binding to epitopes in the C-terminal region of the soluble protein spanning the HR2 region (i.e., multiple phage clones with similar inserts). This region may not be highly exposed on the virions or infected cells but is clearly immunogenic in the soluble recombinant spike ectodomain. In addition, the rabbit anti-S1+S2 antibodies bound diverse epitopes spanning the RBD and to a lesser degree to the N-terminal domain (NTD) and the C-terminal region of S1, and the N-terminus of S2, including the fusion peptide (Fig. 2B and Suppl. Table 1). The S1 domain elicited very strong response against the C-terminal region of S1 protein and a diverse antibody repertoire recognizing the NTD and RBD/RBM regions (Fig. 2C and Suppl. Table 1). The recombinant RBD induced high-titer antibodies that were highly focused to the RBD/RBM (Fig. 2E, and Suppl. Table 1). In contrast, the recombinant S2 immunogen after two immunizations in rabbits elicited antibodies primarily targeting the C-terminus of the S2 protein (CD-HR2).

**Figure 2:**
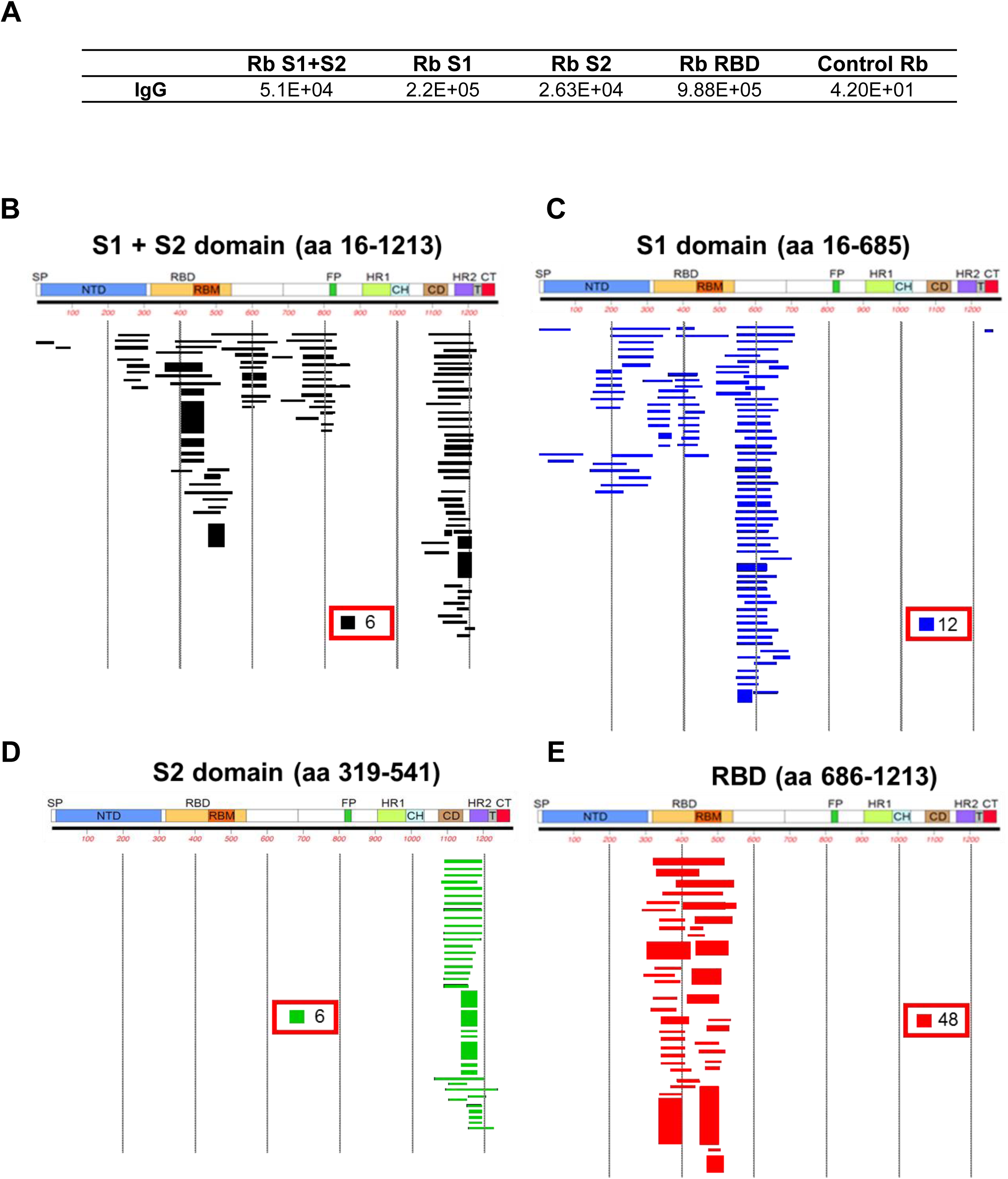
Antibody repertoires generated by different SARS-CoV-2 spike antigens. (A) Number of IgG-bound SARS-CoV-2 GFPDL phage clones using the post-second vaccination rabbit polyclonal sera from the vaccine groups in Fig 1. (B-E) Graphical distribution of representative clones with a frequency of ≥2, obtained after affinity selection, and their alignment to the Spike protein of SARS-CoV-2 are shown for the four vaccine groups: S1+S2 ectodomain (B), S1 (C), S2 domain (D) and S1-Receptor binding domain (RBD) (E). The thickness of each bar represents the frequency of repetitively isolated phage, with the scale shown enclosed in a red box in the respective alignments in each panel. The GFPDL affinity selection data was performed twice. Similar numbers of phage clones and epitope repertoire were observed in both phage display analyses.

All the immunodominant antigenic sites identified by the SARS-CoV-2 GFPDL panning of all 4 immune sera on the spike sequence are shown in Suppl. Fig. 4. Alignment of the sequence with other coronaviruses shows that some of the antigenic sites are >70% conserved among several coronavirus strains isolated from humans and bats, especially those located in the S2 domain (Suppl. Table 1). Structural depiction of these antigenic sites on the SARS-CoV-2 spike (Suppl. Fig. 5; in blue on PDB#6VSB), demonstrated that most of these antigenic sites identified in the current study are surface exposed on the native prefusion spike(*2*).

## DISCUSSION

In this study, we performed an in-depth evaluation of antibody response generated by various SARS-CoV-2 spike antigens that are similar to the vaccine antigens being used in clinical development(*5, 16, 17*). Bioinformatics approach previously identified 279 potential B-cell epitopes and 48 potential T cell epitopes in the Spike glycoproteins of SARS-CoV viruses, based on human antibody responses to the SARS-CoV-1 infection and the corresponding epitopes in SARS-CoV-2 spike (Grifoni et al. Table 4) (*18*). We compared the predominant antigenic sites identified by antibodies in our study generated by different spike antigens with the B cell epitopes predicted by Grifoni *et al*.(*18*). Four of the predicted B epitopes overlapped with the sequences we identified in our GFPDL analysis: aa 287-317 in NTD-RBD overlaps with our antigenic site aa 298-363 which is 77% homologous between SARS-CoV-1 and SARS-CoV-2; aa 524-598 and aa 601-640, in the C-terminus of S1 overlap with our antigenic site containing aa 548-632 (78.8 % conservation between SARS-CoV-1 and SARS-CoV-2); aa 802-819 in the S2 domain/FP overlaps with our antigenic site aa 768-828 (83% conserved between SARS-CoV-1 and SARS-CoV-2) (Fig. S4 and Suppl. Table 1). The other epitopes identified in our study cover less conserved sequences between the two SARS-CoV viruses that are unique to the SARS-CoV-2 spike and were not identified in the *in-silico* approach by Grifoni et al.

Surprisingly, the S2 domain doesn’t appear to elicit as many neutralizing antibodies as RBD or S1. Although S2 contains the fusion peptide, it does not appear to be as immunogenic, compared with S1 or RBD, in generating binding antibodies to the intact spike (S1+S2) ectodomain, as observed in both IgG ELISA and SPR. Even though we characterized the purified proteins in various assays, there is a possibility that the structure of the antigens used in the study is different from the corresponding authentic spike protein on the surface of SARS-CoV-2 virion particle.

One unexpected finding in this study was the higher affinity of antibodies elicited by the RBD compared with the other spike antigens (S1+S2 ectodomain, S1 and S2 domains). In earlier studies, with vaccines against H7N9 avian influenza we found important correlation between antibody affinity against the hemagglutinin HA1 globular domain and control of virus loads after challenge of ferrets with H7N9 (*19*). In study of patients recovering from Zika virus (ZIKV) infections, their antibody affinity against ZIKV E-DIII correlated with lower clinical scores(*20*). In a large randomised clinical trial of IVIG hyper-enriched for influenza virus antibodies (hIVIG), in adults hospitalised with confirmed influenza A or B infections, a statistically significant virological benefit and clinical benefit for patients infected with B strains, directly correlated with stronger antibody affinities of the hIVIG for circulating B strains (*14*). In a recent longitudinal study of Ebola virus disease survivor, affinity maturation to Ebola virus GP was associated with a rapid decline in viral replication and illness severity in this patient (*13*). Thus, vaccines that can elicit high affinity antibodies may have a significant advantage for *in-vivo* clinical outcome of SARS-CoV-2 infection and contribute to amelioration of disease in infected individuals. Therefore, in addition to measurements of antibody-binding titers and virus neutralization, this and the previous studies demonstrate the importance of assessments of antibody affinity maturation during SARS-CoV-2 vaccine trials.

In summary, our study highlights the need to perform comprehensive analysis of immune response generated following vaccination or SARS-CoV-2 infection to identify biomarkers of protective immunity. In-depth understanding of quantitative and qualitative aspects of immune responses generated by different spike protein vaccine antigens could aid the development and evaluation of effective SARS-CoV-2 therapeutics and vaccines.

## ACKNOWLEDGEMENTS

We thank Keith Peden and Marina Zaitseva for their insightful review of the manuscript.

## Funding

The antibody characterization work described in this manuscript was supported by FDA intramural grant funds. The funders had no role in study design, data collection and analysis, decision to publish, or preparation of the manuscript.

The content of this publication does not necessarily reflect the views or policies of the Department of Health and Human Services, nor does mention of trade names, commercial products, or organizations imply endorsement by the U.S. Government.

## AUTHOR CONTRIBUTIONS

**Designed research:** S.K.

**Performed research:** S.R., J. T., E.C., L.K., G. G., S.L., T.W., and S.K.

**Conducted Animal study:** L.K., G. G., and S.K

**SARS-CoV-2 neutralization assays:** S.L., and T.W.

**Contributed to Writing:** H.G. and S.K.

## Materials & Correspondence

Correspondence and material requests should be addressed to the corresponding author (S.K.).

## Ethics Statement

All animal experiments were approved by the U.S. FDA Institutional Animal Care and Use Committee (IACUC) under Protocol #2008-10. The animal care and use protocol meets National Institutes of Health guidelines.

## SUPPLEMENTAL INFORMATION

**Supplementary Figure 1:**
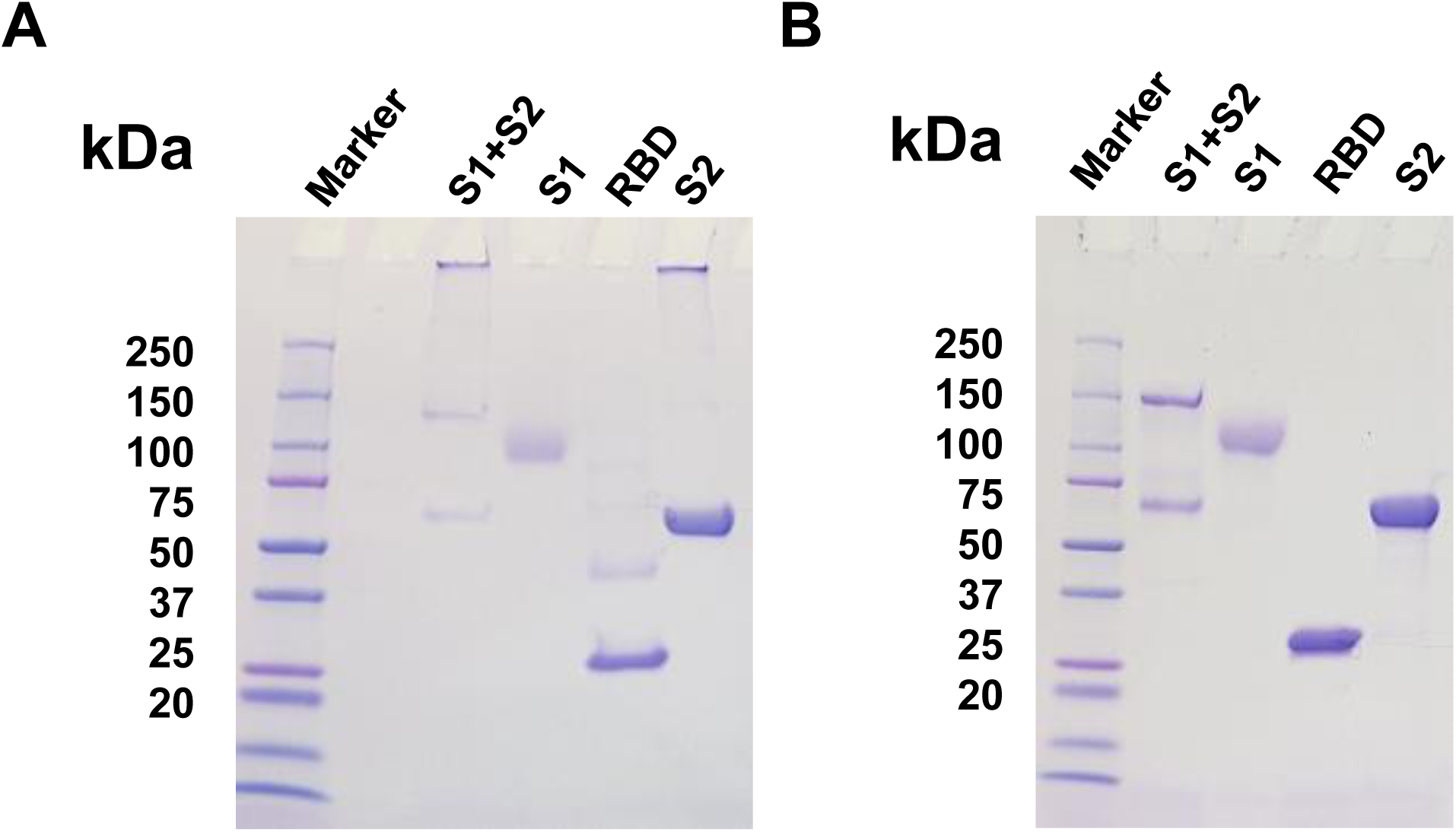
Purified SARS-CoV-2 proteins analyzed by SDS-PAGE and under reducing and non-reducing conditions. Related to figure 1. 2 µg of purified proteins was run in SDS-PAGE under non-reducing (A) and reducing (B) conditions. The gels were stained with Coomassie blue.

**Supplementary Figure 2:**
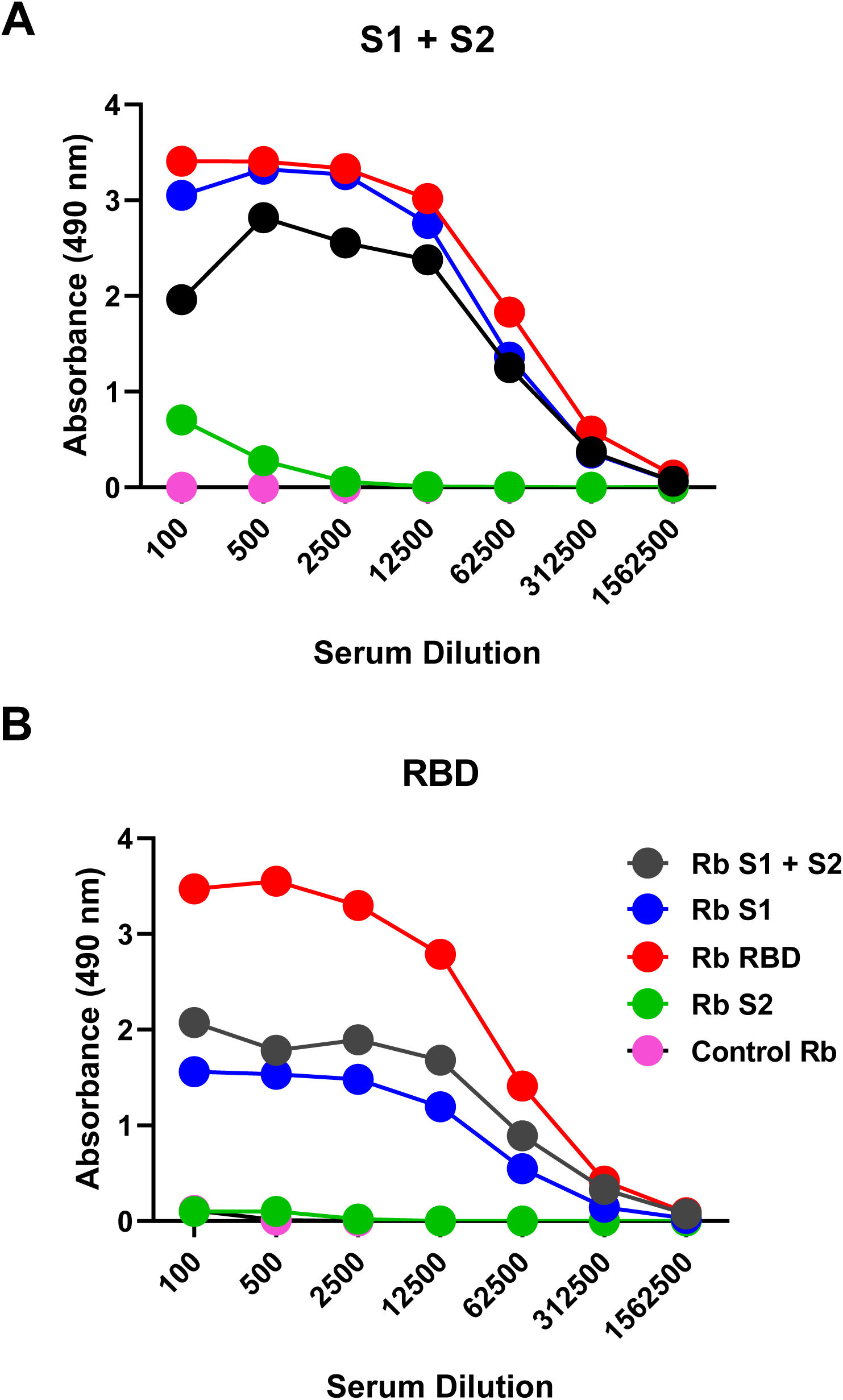
Anti-Spike reactivity of post-vaccination rabbit sera in ELISA. Related to figure 1. Post-vaccination rabbit sera following two immunizations with different SARS-CoV-2 spike vaccine antigens (S1+S2; black, S1; blue, RBD; red, S2; green and pre-vaccination control; pink) in ELISA. Average antibody binding to recombinant Spike (S1+S2) ectodomain (A) and S1-RBD (B) is shown in ELISA using HRP-conjugated goat anti-rabbit IgG specific antibody.

**Supplementary Figure 3.**
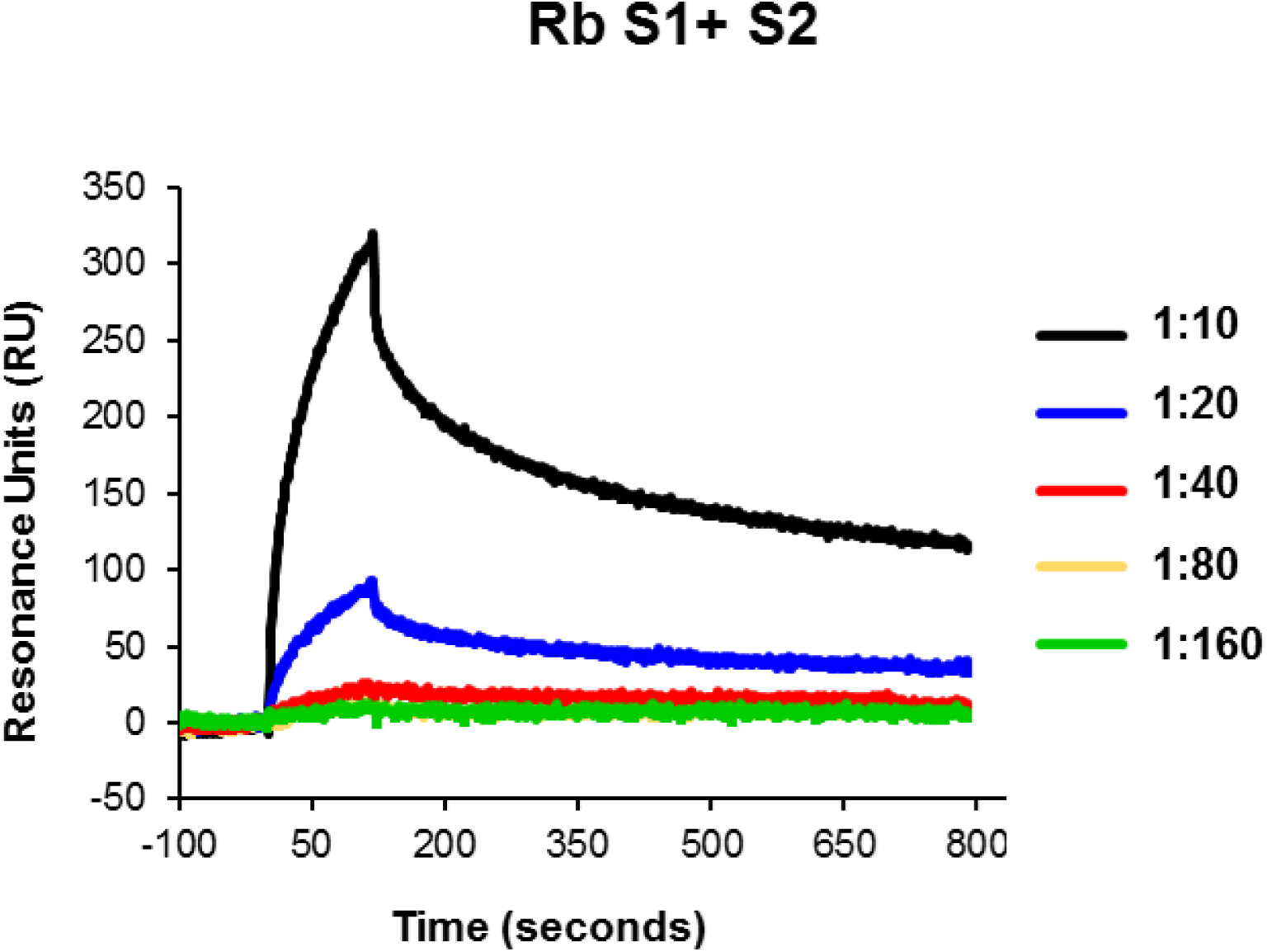
Steady-state equilibrium analysis of serum antibodies binding by SPR. Related to figure 1. Serial dilutions of post-2^nd^ vaccination rabbit antiserum against SARS-CoV-2 Spike (S1+S2 ectodomain) were injected simultaneously onto SARS-CoV-1 S1+S2 captured on a Ni-NTA sensor chip and on a surface free of protein (used as a blank). Binding responses from the protein surface were corrected for the response from the mock surface and for responses from a separate, buffer only injection. Unvaccinated Rabbit control sample at 10-fold dilution did not show any binding in SPR.

**Supplementary Figure 4.**
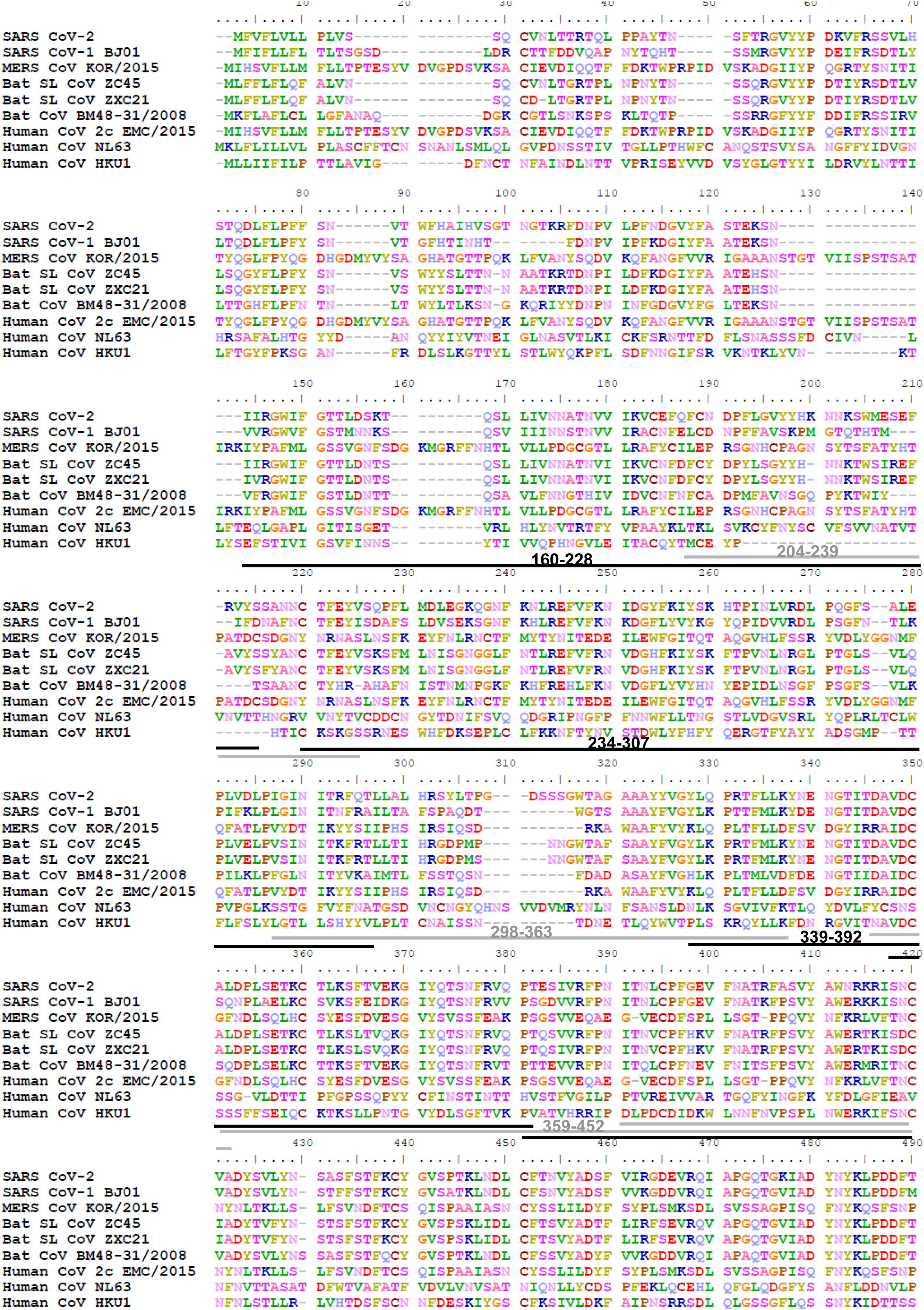
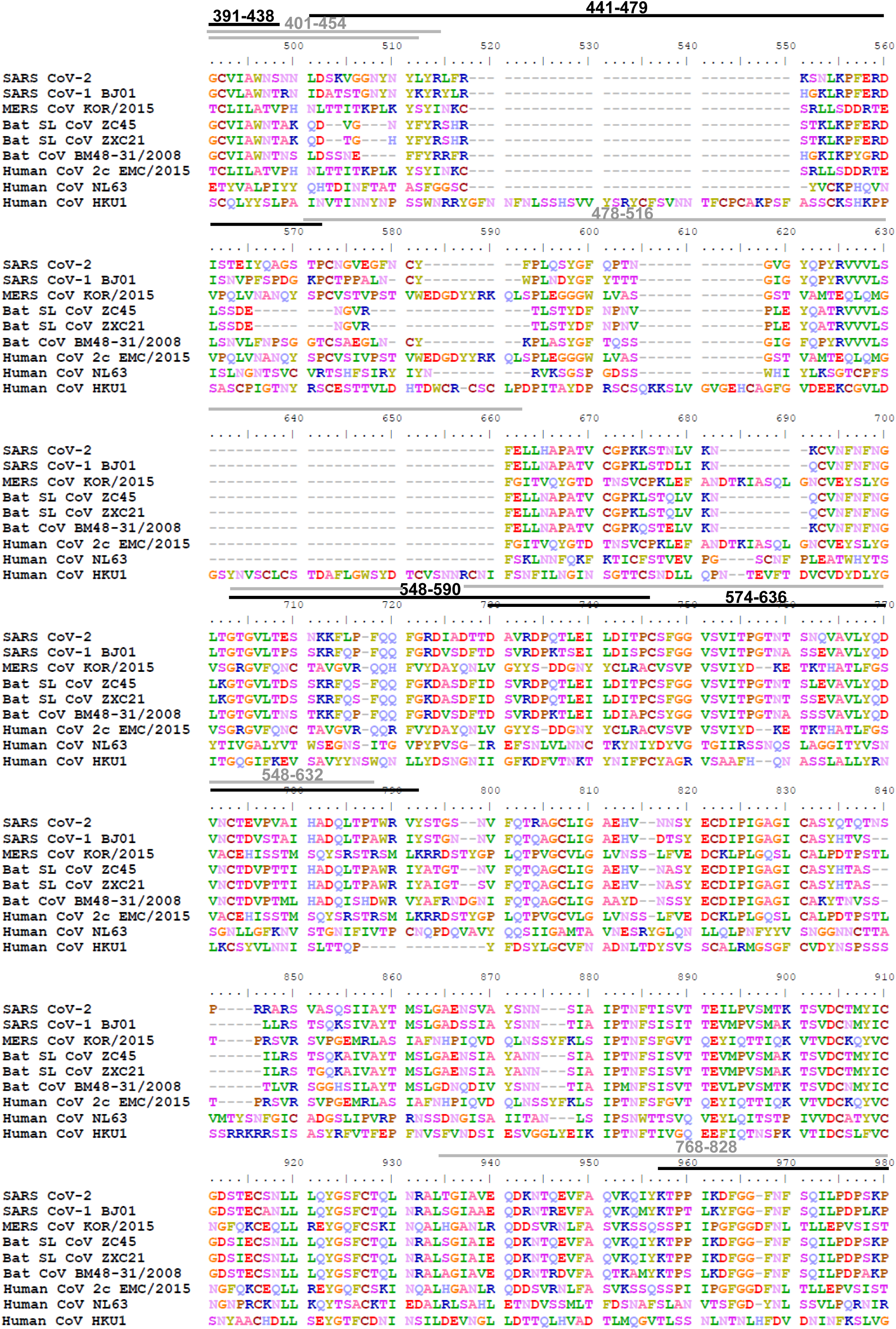
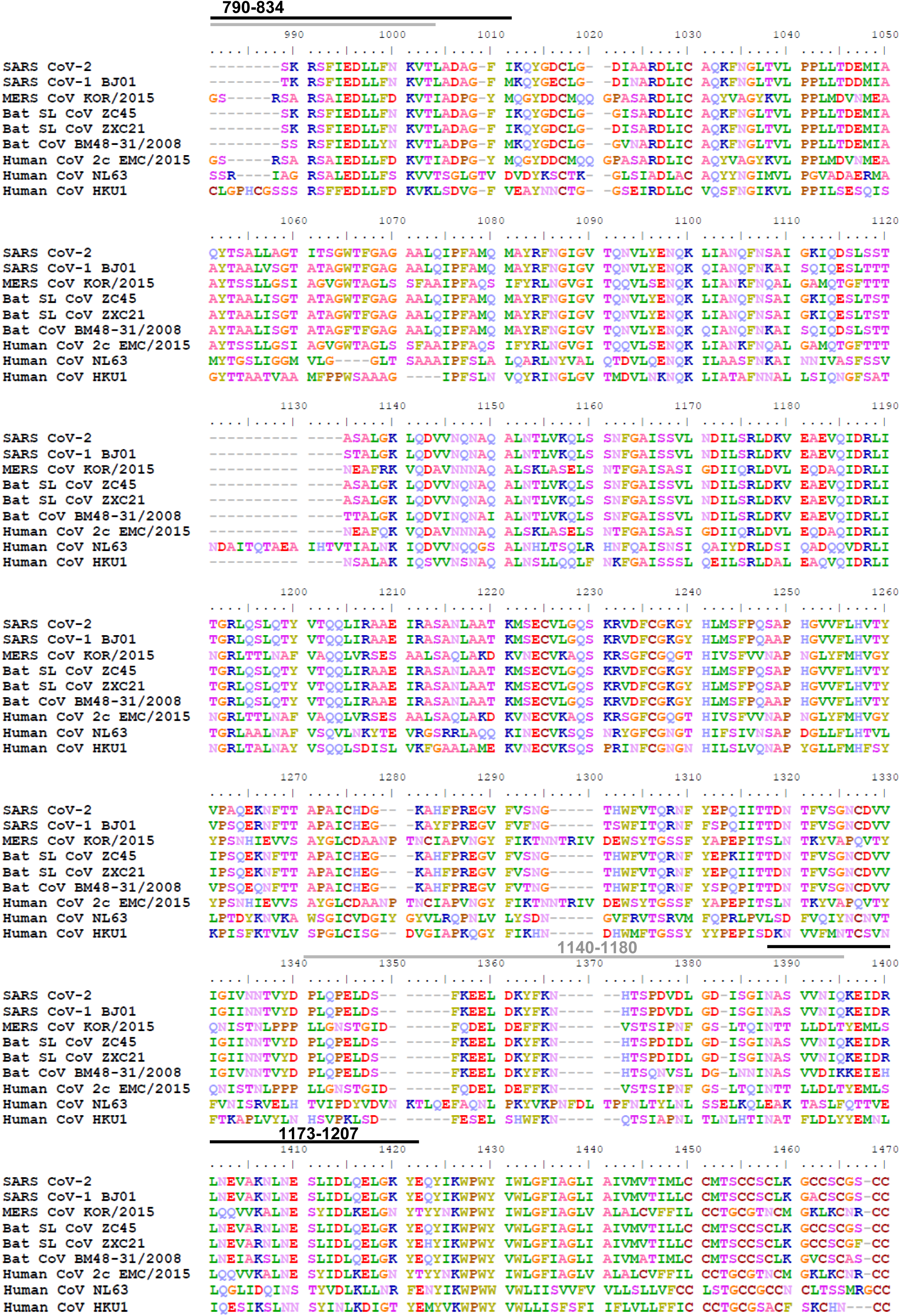
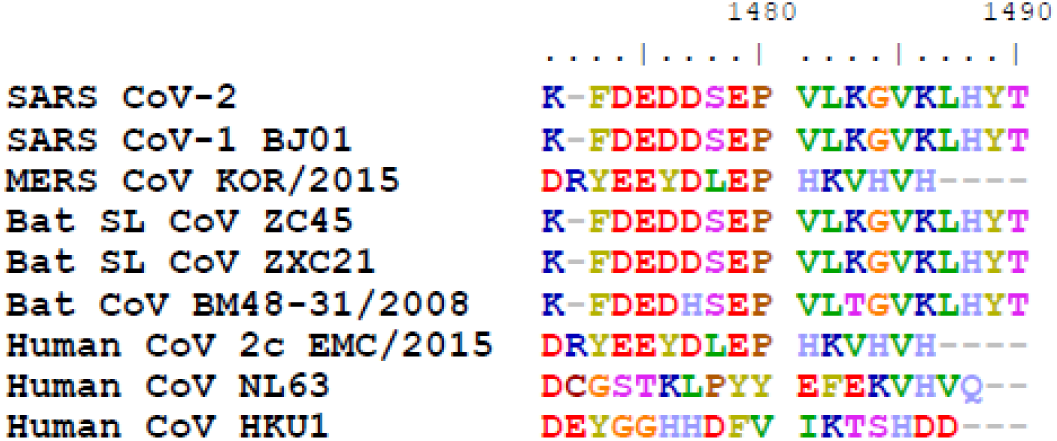
Sequence alignment of Spike protein from diverse CoV strains. Related to figures 1 & 2. An alignment of the spike proteins of SARS-CoV-2 (Genbank#MN908947), SARS-CoV-1 BJ01 strain (Genbank#AAP30030.1), MERS CoV KOR/KNIH/2015(Genbank#AKN11075.1), Bat SARS-like CoV ZC45 (Genbank#AVP78031.1), Bat SARS-like CoV ZXC21 (Genban#AVP78042.1), Bat CoV BM48-31/BGR/2008 (Genbank#ADK66841.1), Human CoV 2c EMC/2012 (Genbank# AFS88936.1), Human CoV NL63 (NCBI#YP_003767.1), and Human CoV HKU1 (NCBI#YP_173238.1) was performed using Clustal W multiple alignment application. Various domains of the spike protein are the S1 subunit (AA 1-685), RBD (AA 319-541), FP (816-834) and S2 (686-1273) subunits. The SARS-CoV-2 antigenic regions/sites discovered in this study using the post-vaccination antibodies with different SARS-CoV-2 vaccine antigens are depicted above the SARS-CoV-2 spike protein sequence in alternating black and grey lines with the corresponding AA residues for visualization.

**Supplementary Figure 5.**
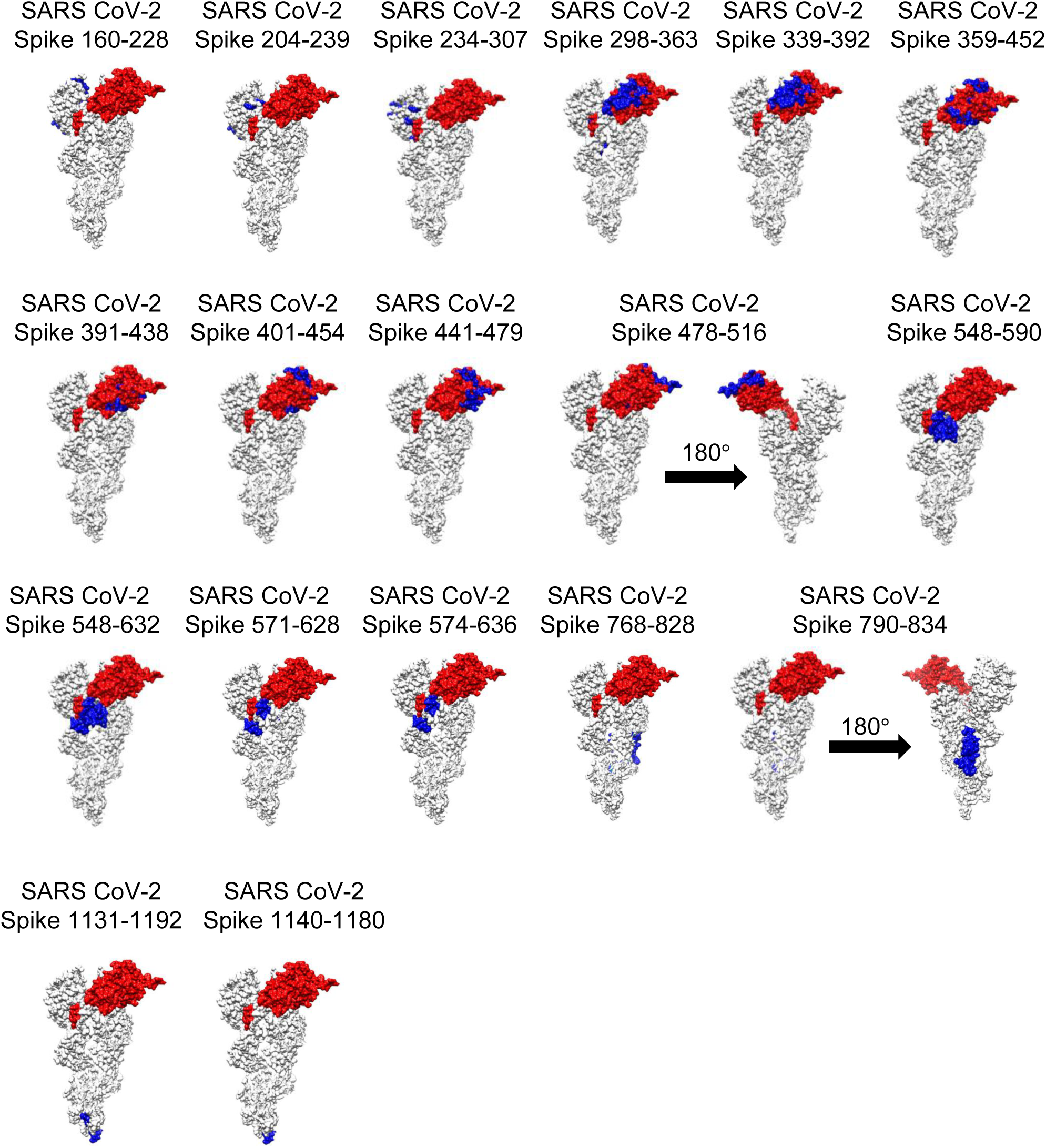
Structural representation of antigenic sites identified in SARS-CoV-2 using GFDPL. Related to figure 2. Antigenic sites identified using GFPDL have been depicted in blue on surface structures of a monomer of PDB#6VSB (Wrapp et al., 2020) with a single receptor-binding domain (RBD) in the up conformation, wherever available using UCSF Chimera software. The RBD region is shaded in red (residues 319-541) on every structure. Those structures (SARS CoV-2 Spike 478-516 and 790-834), whose sites were not visible on the side depicted by flipping the structure by 180°.

**Table S1:** Sequence conservation of Antigenic regions/sites among different CoV strains.

| AA | Similarity of SARS CoV 2 antigenic sites to other CoV strains (%) |  |  |  |  |  |  |  |  |
| --- | --- | --- | --- | --- | --- | --- | --- | --- | --- |
|  | SARS CoV-2 | SARS CoV-1 | MERS | Bat SL CoV ZC45 | Bat SL CoV ZXC21 | Bat CoV BM48 | Human CoV 2c | Human NL63 | Human HKU1 |
| 160-228 | 100 | 53.6 | 9.7 | 62.3 | 60.8 | 39.1 | 9.7 | 12.6 | 5.7 |
| 204-239 | 100 | 55.5 | 10.5 | 63.8 | 63.8 | 44.4 | 10.5 | 21 | 8.1 |
| 234-307 | 100 | 52.7 | 32.4 | 74.3 | 71.6 | 48.6 | 32.4 | 6.4 | 27 |
| 298-363 | 100 | 77.2 | 30.3 | 80.3 | 80.3 | 78.7 | 30.3 | 7.5 | 30.3 |
| 339-392 | 100 | 87 | 24 | 75.9 | 75.9 | 83.6 | 24 | 3.6 | 27 |
| 359-452 | 100 | 80.8 | 20.2 | 74.4 | 72.3 | 76.8 | 20.2 | 5.2 | 20.2 |
| 391-438 | 100 | 83.3 | 20.8 | 83.3 | 83.3 | 81.2 | 20.8 | 8.3 | 22.9 |
| 401-454 | 100 | 74 | 16.6 | 72.2 | 68.5 | 70.3 | 16.6 | 5.5 | 12.9 |
| 441-479 | 100 | 46.1 | 10.2 | 41 | 35.8 | 30.7 | 10.2 | 10.2 | 2.7 |
| 478-516 | 100 | 61.5 | 14.2 | 43.5 | 43.5 | 56.4 | 14.2 | 12.5 | 8.7 |
| 548-590 | 100 | 74.4 | 16.2 | 74.4 | 74.4 | 79 | 16.2 | 4.6 | 22.7 |
| 548-632 | 100 | 78.8 | 17.6 | 80 | 80 | 75.2 | 17.6 | 4.7 | 22 |
| 574-636 | 100 | 80.9 | 14.2 | 87.3 | 87.3 | 76.1 | 14.2 | 3.1 | 22.2 |
| 768-828 | 100 | 83.6 | 41.5 | 96.7 | 96.7 | 80.3 | 41.5 | 24.6 | 24.2 |
| 790-834 | 100 | 86.6 | 46.9 | 100 | 100 | 84.4 | 46.9 | 32 | 31.4 |
| 1140-1180 | 100 | 100 | 26.1 | 97.5 | 97.5 | 75.6 | 26.1 | 14.5 | 33.3 |
| 1173-1207 | 100 | 100 | 45.7 | 97.1 | 97.1 | 82.8 | 45.7 | 20 | 37.1 |

\* Percent sequence conservation of GFPDL identified antigenic sites in the SARS CoV-2 spike protein (Genbank#MN908947), with CoV Spike proteins of SARS CoV-1 BJ01 strain (Genbank#AAP30030.1), MERS CoV KOR/KNIH/2015(Genbank#AKN11075.1), Bat SARS-like CoV ZC45 (Genbank#AVP78031.1), Bat SARS-like CoV ZXC21 (Genban#AVP78042.1), Bat CoV BM48-31/BGR/2008 (Genbank#ADK66841.1), Human CoV 2c EMC/2012 (Genbank# AFS88936.1), Human CoV NL63 (NCBI#YP\_003767.1), and Human CoV HKU1 (NCBI#YP\_173238.1) was calculated using Sequence Identity Matrix function in BioEdit.

## METHODS

### Recombinant CoV Proteins

Recombinant SARS-CoV-2 proteins were purchased from Sino Biologicals (S1+S2 ectodomain; 40589-V08B1, S1; 40591-V08H, RBD; 40592-V08H or S2; 40590-V08B). Recombinant purified proteins used in the study were either produced in HEK 293 mammalian cells (S1 and RBD) or insect cells (S1+S2 ectodomain and S2 domain).

### Rabbit immunization Studies

Female New Zealand white rabbits (Charles River labs) were immunized twice intra-muscularly at 14-days interval with 50 μg of purified proteins mixed with Emulsigen Adjuvant. Sera were collected before (pre-vaccination) and after 1^st^ and 2^nd^ vaccination and analyzed for binding antibodies in ELISA, SPR, neutralization assay and GFPDL analysis.

### ELISA

96 well Immulon plates were coated with 100 ng/100 µL of recombinant spike protein and protein domains in PBS overnight at 4°C. Starting at a 1:100 dilution, serum samples were serially diluted 1:5 and applied to the protein-coated plate in 10 µL for 1 hr at ambient temperature. Serum samples were assayed in duplicate. Naïve serum samples were assayed along with the experimental samples. After three washes with PBS/0.05% Tween 20, bound antibodies were detected with a donkey anti-rabbit IgG Fc-specific HRP-conjugated antibody (Jackson Immuno Research) After 1hr, plates were washed as before and OPD was added for 10min. Absorbance was measured at 492 nm. End titer was determined as 2-fold above the average of the absorbance values of the naïve serum samples. The end titer is reported as the last serum dilution that was above this cutoff.

### Antibody binding kinetics of post-vaccination sera to recombinant SARS-CoV-2 proteins by Surface Plasmon Resonance (SPR)

Steady-state equilibrium binding of post-vaccination rabbit polyclonal serum was monitored at 25°C using a ProteOn surface plasmon resonance (BioRad). The purified recombinant Spike proteins were captured to a Ni-NTA sensor chip with 200 resonance units (RU) in the test flow channels. The native functional activity of the Spike proteins was determined by binding to the 5 µg/mL human ACE2 protein.

For serum analysis, the protein density on the chip was optimized such as to measure monovalent interactions independent of the antibody isotype. Serially diluted (10-, 20-, 40-, 80-, and 160-fold of freshly prepared sera were injected at a flow rate of 50 µl/min (120 sec contact duration) for association, and disassociation was performed over a 600-second interval. Responses from the protein surface were corrected for the response from a mock surface and for responses from a buffer-only injection. SPR was performed with serially diluted serum of each animal in this study. Antibody isotype analysis for the SARS-CoV-2 spike protein bound antibodies in the polyclonal serum was performed using SPR. Total antibody binding was calculated with BioRad ProteOn manager software (version 3.1). All SPR experiments were performed twice and the researchers performing the assay were blinded to sample identity. In these optimized SPR conditions, the variation for each sample in duplicate SPR runs was <5%. The maximum resonance units (Max RU) data shown in the figures was the RU signal for the 10-fold diluted serum sample. Antibody off-rate constants, which describe the fraction of antigen–antibody complexes that decay per second, are determined directly from the serum/ sample interaction with SARS CoV-2 spike ectodomain (S1+S2), S1, S2, and RBD using SPR in the dissociation phase only for the sensorgrams with Max RU in the range of 20–100 RU and calculated using the BioRad ProteOn manager software for the heterogeneous sample model as described before(*11*). Off-rate constants were determined from two independent SPR runs.

### SARS-CoV-2 pseudovirus production and neutralization assay

Human codon-optimized cDNA encoding SARS-CoV-2 S glycoprotein (NC_045512) was synthesized by GenScript and cloned into eukaryotic cell expression vector pcDNA 3.1 between the BamH*I* and Xho*I* sites. Pseudovirions were produced by co-transfection Lenti‐X 293T cells with pMLV-gag-pol, pFBluc, and pcDNA 3.1 SARS-CoV-2 S using Lipofectamine 3000. The supernatants were harvested at 48h and 72h post transfection and filtered through 0.45-mm membranes.

For neutralization assay, 50 µL of SARS-CoV-2 S pseudovirions were pre-incubated with an equal volume of medium containing serum at varying dilutions at room temperature for 1 h, then virus-antibody mixtures were added to Vero E6 cells in a 96-well plate. After a 3 h incubation, the inoculum was replaced with fresh medium. Cells were lysed 48 h later, and luciferase activity was measured using luciferin-containing substrate.

### Gene Fragment Phage Display Library (GFPDL) construction

SARS-CoV-2 spike gene was chemically synthesized and used for cloning and construction of phage display libraries. A gIII display-based phage vector, fSK-9-3, was used where the desired polypeptide can be displayed on the surface of the phage as a gIII-fusion protein. Purified DNA containing spike gene was digested with *DNase I* to obtain gene fragments of 100-1000 bp size range and used for GFPDL construction as described previously (*6-8*). The phage libraries constructed from the SARS-CoV-2 spike gene display viral protein segments ranging in size from 30 to 350 amino acids, as fusion protein on the surface of bacteriophage.

### Affinity selection of SARS-CoV-2 GFPDL phages with polyclonal rabbit serum

Prior to panning of GFPDL with polyclonal serum antibodies, serum components that could non-specifically interact with phage proteins were removed by incubation with UV-killed M13K07 phage-coated petri dishes (*8*). Equal volumes of each post-vaccination rabbit serum were used for GFPDL panning. GFPDL affinity selection was carried out in-solution with protein A/G (IgG) specific affinity resin as previously described (*6, 7, 9*) Briefly, the individual rabbit serum was incubated with the GFPDL and the protein A/G resin, the unbound phages were removed by PBST (PBS containing 0.1 % Tween-20) wash followed by washes with PBS. Bound phages were eluted by addition of 0.1 N Gly-HCl pH 2.2 and neutralized by adding 8 µL of 2 M Tris solution per 100 µL eluate. After panning, antibody-bound phage clones were amplified, the inserts were sequenced, and the sequences were aligned to the SARS-CoV-2 spike gene, to define the fine epitope specificity in the post-vaccination rabbit sera. The GFPDL affinity selection data was performed blindly in a blinded fashion. Similar numbers of bound phage clones and epitope repertoire were observed in the two GFPDL panning.

### Data Availability

The datasets generated during and/or analyzed during the current study are available from the corresponding author on reasonable request.

